# Infection dynamics of co-transmitted reproductive symbionts are mediated by sex, tissue, and development

**DOI:** 10.1101/2022.03.25.485896

**Authors:** Megan W Jones, Laura C Fricke, Cody J Thorpe, Lauren O Vander Esch, Amelia RI Lindsey

## Abstract

One of the most prevalent intracellular infections on earth is with *Wolbachia*: a bacterium in the Rickettsiales that infects a range of insects, crustaceans, chelicerates, and nematodes. *Wolbachia* is maternally transmitted to offspring and has profound effects on the reproduction and physiology of its hosts, which can result in reproductive isolation, altered vectorial capacity, mitochondrial sweeps, and even host speciation. Some populations stably harbor multiple *Wolbachia* strains, which can further contribute to reproductive isolation and altered host physiology. However, almost nothing is known about the requirements for multiple intracellular microbes to be stably maintained across generations while they likely compete for space and resources. Here we use a coinfection of two *Wolbachia* strains (“*w*Ha” and “*w*No”) in *Drosophila simulans* to define the infection and transmission dynamics of an evolutionarily stable double infection. We find that a combination of sex, tissue, and host development contribute to the infection dynamics of the two microbes and that these infections exhibit a degree of niche partitioning across host tissues. *w*Ha is present at a significantly higher titer than *w*No in most tissues and developmental stages, but *w*No is uniquely dominant in ovaries. Unexpectedly, the ratio of *w*Ha to *w*No in embryos does not reflect those observed in the ovaries, indicative of strain-specific transmission dynamics. Understanding how *Wolbachia* strains interact to establish and maintain stable infections has important implications for the development and effective implementation of *Wolbachia*-based vector biocontrol strategies, as well as more broadly defining how cooperation and conflict shape intracellular communities.

**IMPORTANCE:** *Wolbachia* are maternally transmitted intracellular bacteria that manipulate the reproduction and physiology of arthropods, resulting in drastic effects on the fitness, evolution, and even speciation of their hosts. Some hosts naturally harbor multiple strains of *Wolbachia* that are stably transmitted across generations, but almost nothing is known about the factors that limit or promote these co-infections which can have profound effects on the host’s biology and evolution, and are under consideration as an insect-management tool. Here we define the infection dynamics of a known stably transmitted double infection in *Drosophila simulans* with an eye towards understanding the patterns of infection that might facilitate compatibility between the two microbes. We find that a combination of sex, tissue, and development all contribute how the coinfection establishes.

## INTRODUCTION

Eukaryotic cells are home to a diversity of intracellular microbes including mitochondria, plastids, symbionts, and pathogens, many of which are vertically inherited via the maternal germline. The community and interactions between intracellular microbes are associated with diverse effects on host physiology and health. Despite the importance of the intracellular community, little is known about the factors that promote, inhibit, or regulate the establishment and transmission of multiple, coinfecting, intracellular microbes.

Arthropods are particularly rich in examples of such infections. It is estimated that more than half of arthropods have at least one heritable bacterial symbiont, and ∼12% have two or more of these infections (1, 2).The most common of these is an alpha-proteobacterium, *Wolbachia*, a close relative of the intracellular human pathogens *Anaplasma, Rickettsia*, and *Ehrlichia* (3). Unlike their close relatives, *Wolbachia* inhabit the cells of arthropods and nematodes, are primarily vertically transmitted via the maternal germline, and alter host physiology and reproduction to facilitate spread through a population (4, 5). Some arthropods stably harbor multiple co-infecting *Wolbachia* strains (6-10), resulting in drastic effects on host fitness, gene flow between populations, horizontal transfer between *Wolbachia*, and even host speciation (8, 10-15). Not only are *Wolbachia* coinfections significant for evolution of both the microbes and the arthropod host, but the increasing interest in establishing secondary *Wolbachia* infections for use in insect control programs necessitates a mechanistic investigation of these intracellular inhabitants (16-18). Previous successes in *Wolbachia*-mediated vector control were more easily attainable because key vector species such as *Aedes aegypti* so happened to naturally lack *Wolbachia* (19, 20). However, many other pest and vector species are already infected with resident *Wolbachia* strains, and establishment of a secondary infection is a potential avenue for control methods (17, 18, 21). Furthermore, pathogens and symbionts in related systems are rarely in complete isolation and the intracellular interactions between symbiotic microbes, pathogenic microbes, mitochondria, and viruses can all contribute to altered host physiology, vector competence, and/or clinical progression of disease (22-27).

While very little is known about the infection dynamics of co-occurring *Wolbachia*, there are several shared characteristics across many of the naturally occurring *Wolbachia* coinfections, indicating there may be shared mechanisms and selective pressures at play. For example, in *Aedes albopictus* infected with *w*AlbA and *w*AlbB *Wolbachia* strains (10), *Nasonia vitripennis* (with *w*VitA and *w*VitB (7)), *Dactylopius coccus* (with *w*DacA and *w*DacB (28)), and *Drosophila simulans* (with *w*Ha and *w*No (12)), each insect has one *Wolbachia* strain from supergroup A and one from supergroup B: perhaps indicating that more divergent strains are more compatible in a co-infection, maybe as a result of niche partitioning. In support of this idea, a recent study describing an artificially generated triple infection of *Wolbachia* strains in *Aedes albopictus* showed there was strong competition between *Wolbachia* from the same supergroup, but not between *Wolbachia* from different supergroups (29). There are other examples of artificially generated multiple infections, but the outcomes are highly variable: sometimes the infection destabilizes and is quickly lost, other times it is stable across many generations (30-35). Ultimately, we do not know which factors facilitate successful establishment and transmission of multiple *Wolbachia* strains within one host matriline.

There is literature that suggests the titers of individual strains are differentially regulated. In *Aedes albopictus* mosquitoes, the native *w*AlbB strain is present at ∼6X the titer of the coinfecting native *w*AlbA strain (9). In *Drosophila simulans*, the *w*Ha and *w*No strains establish at different titers in mono-infection conditions, and these titers depend on the combination of strain identity and host tissue (36, 37). However, studies that investigated these strain-specific dynamics leveraged independent fly genetic backgrounds that carried either the *w*Ha strain or *w*No strain, which confounds our interpretation of coinfection dynamics (12, 36-38).

Broadly, there is evidence for both (1) host control over the titer of individual *Wolbachia* strains, and/or (2) the presence of a coinfecting strain contributing to the regulation of *Wolbachia* density (39, 40). However, we have limited knowledge of (1) how coinfecting strains might establish across host tissues and developmental stages, (2) if coinfecting strains facilitate each other’s transmission, (3) if strains evolved to occupy unique niches within the host, (4) if strains go through different severities of population bottleneck from ovary to oocyte, (5) if there are combinatorial effects of the coinfection on host physiology, and ultimately, (6) the host and microbial mechanisms that regulate the maintenance of these coinfections. To begin to investigate these questions, we explore infection and transmission dynamics of multiple vertically inherited intracellular symbionts in a *Drosophila simulans* model which naturally harbors a stable coinfection of two *Wolbachia* strains: *w*Ha and *w*No.

## METHODS

### Bioinformatics

Protein sequences from the reference genomes of *w*Ha (GCF_000376605.1) and *w*No (GCF_000376585.1) annotated with PGAP (8, 41) were used to build orthologous groups of *Wolbachia* proteins using ProteinOrtho v5.15 with default parameters (42). Functional annotations were designated with BlastKOALA with (taxonomy group = bacteria) and (database = eukaryotes + prokaryotes) (43). A *Wolbachia* strain phylogeny was reconstructed with FtsZ sequences from A and B supergroup *Wolbachia*, and a D-supergroup *Wolbachia* (*w*Bm) as outgroup (Supplemental Table S1). Amino acid sequences were aligned with MAFFT and a simple Neighbor Joining (NJ) algorithm was used to reconstruct relationships including a JTT substitution model and 100 bootstrap replicates (44). Tree topology was visualized in FigTree v.1.4.4 (https://github.com/rambaut/figtree) prior to annotation in Inkscape v.1.1.2 (https://inkscape.org/).(44)

### Fly husbandry

Fly stocks were maintained on standard Bloomington cornmeal-agar medium (Nutri-fly® Bloomington Formulation) at 25 °C on a 24-hour, 12:12 light:dark cycle under density-controlled conditions and 50% relative humidity. Experiments used the *Drosophila simulans* genome reference line (Cornell Stock Center SKU: 14021-0251.198), originally from Noumea, New Caledonia, which is stably coinfected with the *w*No and *w*Ha *Wolbachia* strains (12). We generated a *Wolbachia*-free stock with antibiotics for use as a negative control. This stock was generated by tetracycline treatment (20 µg/mL in the fly food for three generations), followed by re-inoculation of the gut microbiome by transfer to bottles that previously harbored male flies from the original stock that had fed and defecated on the media for one week (45). Gonad dissections were performed on live anesthetized flies under sterile conditions, and tissues were immediately flash frozen and stored at −80 °C for later processing. Embryo collections and developmental synchronization was performed using timed 2-hour egg-lays in mating cages on grape agar plates streaked with yeast-paste. For developmental time points, single embryos were collected at two and ten hours, and the remaining embryos were transferred to BSDC media after which single flies were collected as L1, L2, and L3 larvae, white-prepupae, red-eye bald pupae, and pharate males and females (less than two hours post emergence).

### *Wolbachia* screening

Infection status of all stocks was regularly screened with a multiplex PCR assay that produces size-specific amplicons for *w*Ha and *w*No (46). This PCR assay was also used in determining strain segregation during the differential curing experiments (see below). In all cases, DNA was extracted from individual flies with the Monarch® Genomic DNA Purification Kit (New England Biolabs), PCR assays were performed with the strain-specific multiplex primers from (46) and Q5® Hot Start High-Fidelity 2X Master Mix (New England Biolabs) in 20 µl reactions, and products were run on a 1% agarose gel, stained post-electrophoresis with GelRed® (Biotium). For samples that screened negative for *Wolbachia*, DNA integrity was confirmed with PCR using general primers that target arthropod 28S (6). All primer sequences are listed in Table 1.

**Table 1.**
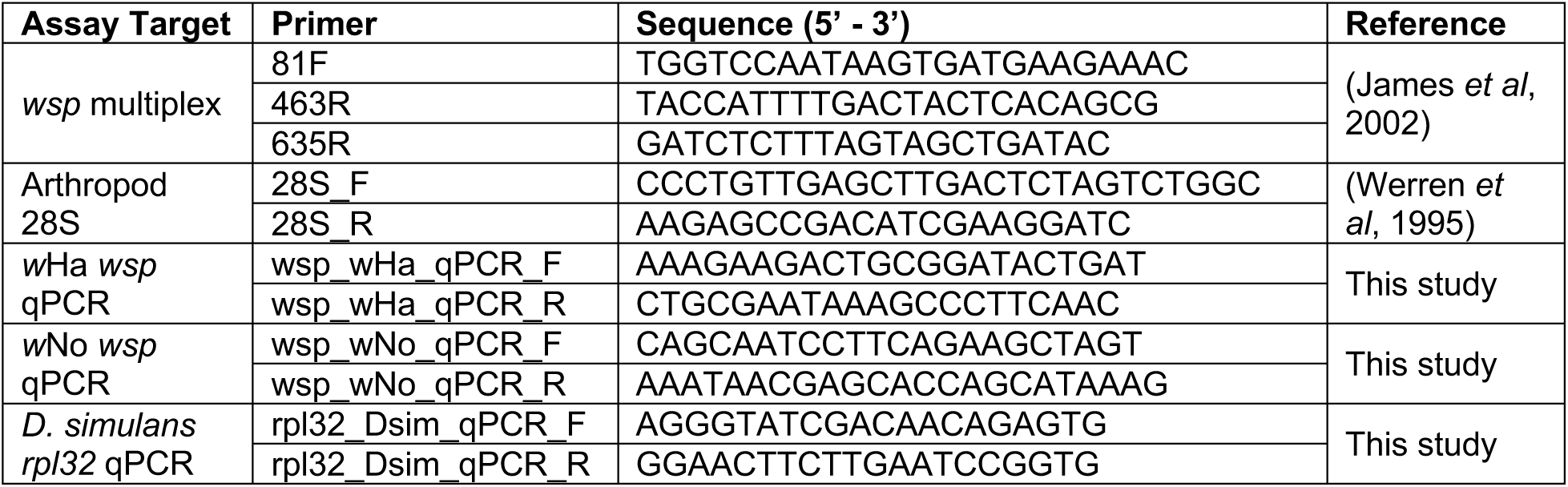
Primer sequences used in this study.

### Strain specific quantitative PCR (qPCR)

To quantify the relative abundance of individual *Wolbachia* strains, we designed *w*Ha- and *w*No-specific qPCR primer sets targeting unique ∼100bp amplicons of the *Wolbachia surface protein* (*wsp*). Assay specificity was verified with Sanger sequencing of amplicons, combined with validation against mono-infected samples generated during differential curing (see above). DNA was extracted from flies/tissues with the Monarch® Genomic DNA Purification Kit (New England Biolabs). Strain specific abundance was assessed with the Luna® Universal qPCR Master Mix (New England Biolabs) following manufacturer’s instructions, and normalization to host genome abundance via amplification of *rpl32*. All reactions were run in technical triplicate alongside a standard curve and negative controls on an QuantStudio™ 3 Real-Time PCR System (Applied Biosystems™). All primer sequences are listed in Table 1.

### Differential curing of *Wolbachia* strains

To disrupt coinfection transmission, we designed a partial heat-cure to reduce *Wolbachia* titers and increase the severity of the bottleneck as *Wolbachia* are deposited in each embryo. Bottles of ∼200 *Drosophila simulans* were kept at 30 °C for four days (or at 25 °C as a control), after which flies were transferred to fresh media under standard rearing conditions (see above) and allowed to oviposit for three days. Offspring (adults <24 hours post eclosion) of the heat-treated mothers were collected and stored in ethanol for further processing.

### Statistics and Data Visualization

All statistics and data visualization were carried out in R version 3.5.0 (47). We used permutational multivariate analysis of variance with the adonis function from the vegan package (48) to assess variation in coinfection titers (a multivariate response) across fly samples using Euclidean distance and 1,000 permutations. Fixed effects were specific to each experimental analysis and included: sex, mating status, and the interaction of the two (Figure 2A), tissue, sex, and the interaction of the two (Figure 2B), or developmental stage (Figure 3). Pairwise comparisons were performed with a Mann-Whitney U test (function “wilcox.test”) followed by Bonferroni Corrections in the case of multiple testing. In the case of the mated vs unmated ovary samples (Figure 4A), we were interested in strain-specific dynamics upon mating, so we assessed variation in strain titers with a two-way ANOVA (function “aov”) including “strain” and “mated status”, along with their interaction, as fixed effects. Correlation between abundance of strains or between abundance in different tissues was assessed with a Spearman’s rank correlation for the data in Figure 2 (function “cor.test”, method= “spearman”). Linear regression was performed with the “lm” function.

## RESULTS

### Coinfecting strains *w*Ha and *w*No share 75% of their coding sequences

To better understand the factors that might facilitate compatibility of two strains we used a suite of bioinformatic approaches to look at phylogenetic and genomic patterns of *Wolbachia* coinfections. Our focal strains, *w*Ha and *w*No (from supergroup A and B, respectively) that coinfect some populations of *Drosophila simulans*, share 858 orthologous groups of proteins, approximately 75% of the coding content of each strain (Figure 1A). The remaining ∼300 proteins in each strain that are not shared are largely hypothetical, unannotated protein sequences, and only 10-15% were assigned a putative function (*w*Ha n = 31/303; *w*No = 44/299). Annotated proteins (i.e., assigned a KEGG KO term) specific to *w*No included 16 transposases, 15 proteins that were related to transcription, DNA repair, or endonuclease activity, and the remaining were largely metabolic in predicted function (Supplemental Table S2). Notably, *w*No encodes for a putative multidrug efflux pump that is not present in *w*Ha. *w*Ha-specific proteins included 15 transposases, three proteins predicted to be involved in transcription or DNA repair, and then a suite of proteins mostly with predicted functions in amino acid transport and metabolism. Interestingly, the *w*Ha strain has two proteins for an addiction module toxin (RelE/StbE family), and a predicted eukaryotic-like golgin-family protein, potentially an effector protein that could interact with host intracellular membranes.

**Figure 1.**
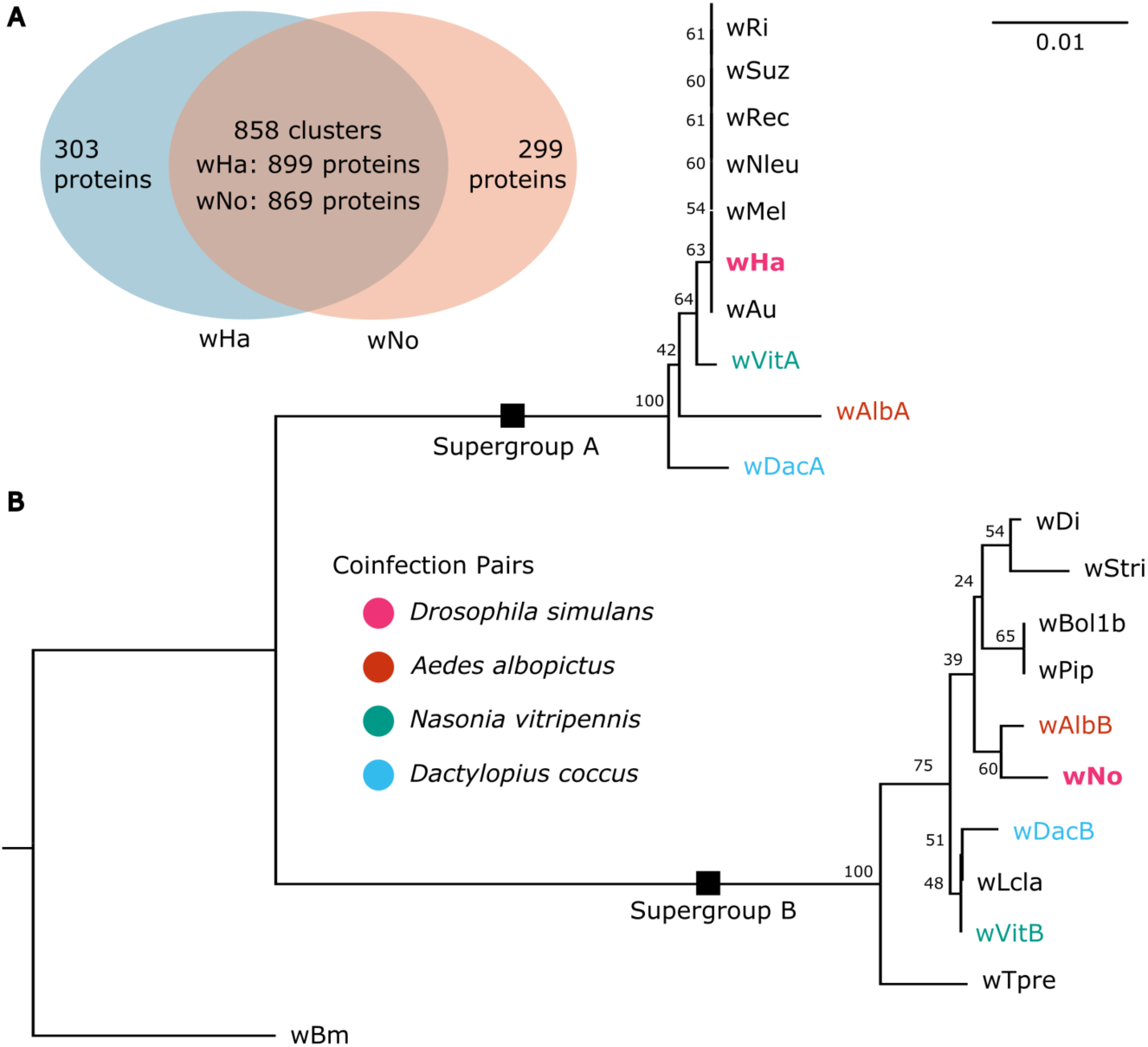
Coinfecting *Wolbachia* strains. **(A)** Shared and unique genes between the focal strains *w*Ha and *w*No that coinfect *Drosophila simulans*. **(B)** Phylogenetic reconstruction of A- and B-supergroup *Wolbachia* based on FtsZ protein sequences, with colors indicating pairs of *Wolbachia* strains that can be found together within a given host. Node labels indicate bootstrap support (n = 100 replicates).

### Strain-specific titers are sex dependent

We assessed the titers of the *w*Ha and *w*No strains in whole body three-day old unmated males and females, and three-day old males and females 24 hours post mating (Figure 2A). There was a significant effect of the interaction between fly sex and mated status (F_1,33_ = 4.076, p = 0.033) as well as a significant effect of sex alone (F_1,33_ = 69.568, p = 0.001), but not of mated status alone (F_1,33_ = 0.488, p = 0.500). This was seen as relatively equal titers of *w*Ha and *w*No in female flies that slightly increased in total abundance upon mating. In contrast, males had drastically reduced titers of *w*No, both relative to *w*No in females, and relative to the coinfecting *w*Ha stain within a male. *w*Ha titers were slightly reduced in males upon mating. Together, these data indicate strong sex-dependent effects on coinfection dynamics.

**Figure 2.**
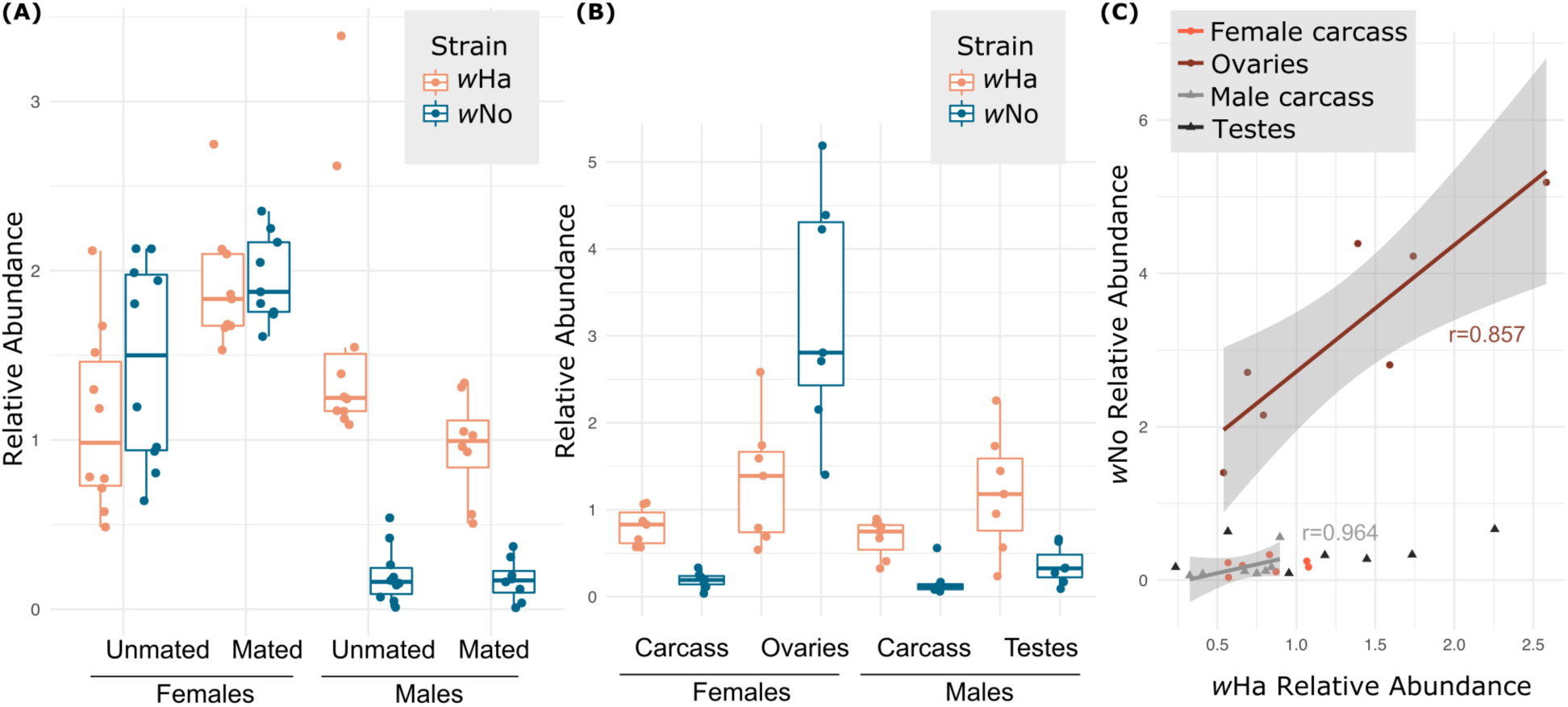
Infection densities of coinfecting *Wolbachia* strains. **(A)** *w*Ha and *w*No titers in whole body mated and unmated males and females. There was a significant effect of the interaction between fly sex and mated status (F_1,33_ = 4.076, p = 0.033) and sex (F_1,33_ = 69.568, p = 0.001) on the coinfection. **(B)** *w*Ha and *w*No titers of gonads and carcasses of unmated males and females. The interaction of sex and tissue significantly affected the coinfection (F_1,27_ = 19.334, p = 0.001), as well as sex alone and tissue alone (F_1,27_ = 19.982, p = 0.001, and, F_1,27_ = 27.147, p = 0.001, respectively). **(C)** Correlation between *w*Ha and *w*No abundance within each sample. Regression lines are shown for ovaries and male carcasses, for which we identified significant correlations in strain-specific abundance (see main text).

### Coinfection dynamics are sex and tissue dependent

A subset of the unmated males and females were dissected prior to DNA extraction resulting in paired gonadal and “carcass” (all remaining tissue) samples for each fly. Strain specific qPCR revealed that the interaction of sex and tissue identity had a significant effect on the abundance of the two strains in the coinfection (F_1,27_ = 19.334, p = 0.001). Additionally, there was significant effect of sex alone, and tissue alone (F_1,27_ = 19.982, p = 0.001, and, F_1,27_ = 27.147, p = 0.001, respectively). In contrast to the relatively equal titers of *w*Ha and *w*No seen in whole female samples (Figure 2A), we found that ovaries were highly enriched for the *w*No strain (Figure 2B). However, in all other sample types (female carcasses, male testes, male carcasses), the *w*Ha strain was significantly more abundant.

We then tested for correlation between the relative abundance of *w*Ha and *w*No within a sample type. We found that in ovaries and male carcasses, there was a significant positive correlation between the abundance of *w*Ha and *w*No (rho = 0.0238, p = 0.8571, and, rho = 0.9643, p =0.0023, respectively). However, in testes and female carcasses, titers of *w*Ha and *w*No were uncorrelated (rho = 0.0714, p = 0.9063, and rho = 0.5357, p = 0.2357, respectively). Next, we asked if there was any correlation in the coinfection between samples that originated from the same fly. We did this in two ways: (1) by comparing the ratio of *w*Ha and *w*No within the gonads, to the same ratio in the carcass, and (2) by comparing the total abundance of *w*Ha and *w*No between gonads and carcass. In both cases, we found no significant relationship between the infection dynamics in the gonads and the carcass (Supplemental Figure S1). In fact, female flies had a very consistent ratio of *w*Ha to *w*No in the ovaries (0.39 +/-0.1) and highly variable *w*Ha:*w*No ratios in the carcass (6.08 +/-4.69). In agreement with the data shown in Figure 2B, the opposite is true in males: the *w*Ha:*w*No ratio is more consistent in the carcass, but highly variable in the testes (Supplemental Fig S1).

### The coinfection is dynamic across development

Given the difference in coinfection between sexes and tissues, we wondered if this was due to differences in transmission of *Wolbachia* to embryos and/or changes across development. To test this, we set up timed egg-lays and collected a developmental series that included seven timepoints across development (from 2-hour old embryos to red-eye-bald pupal stage) as well as newly emerged pharate males and females (Figure 3). Strain-specific qPCR revealed that the coinfection changed significantly across development (Figure 3; F_8,59_ = 2.6682, p = 0.01). Note that juvenile stages were collected without regard to sex, but there were no indications of bimodal distributions which might indicate that juvenile males and females had drastically different patterns of infection. Notably, the pattern of infection in very young embryos did not resemble any of the previously assessed sample types, including the ovaries. Indeed, 2-hour old embryos had more equal titers of *w*Ha and *w*No, unlike the strong *w*No bias in ovaries, and unlike the strong *w*Ha bias in carcasses and testes. By the first larval instar (L1), the coinfection converged on a pattern more similar to the carcass tissue and testes, where *w*Ha titers were much higher than *w*No. This pattern was relatively stable throughout development. In the newly eclosed pharate females there was a significant increase in *w*No titer relative to the pharate males (p = 0.0286) likely indicative of a shift towards the *w*No bias we saw in three-day old female ovaries (Figure 2B).

**Figure 3.**
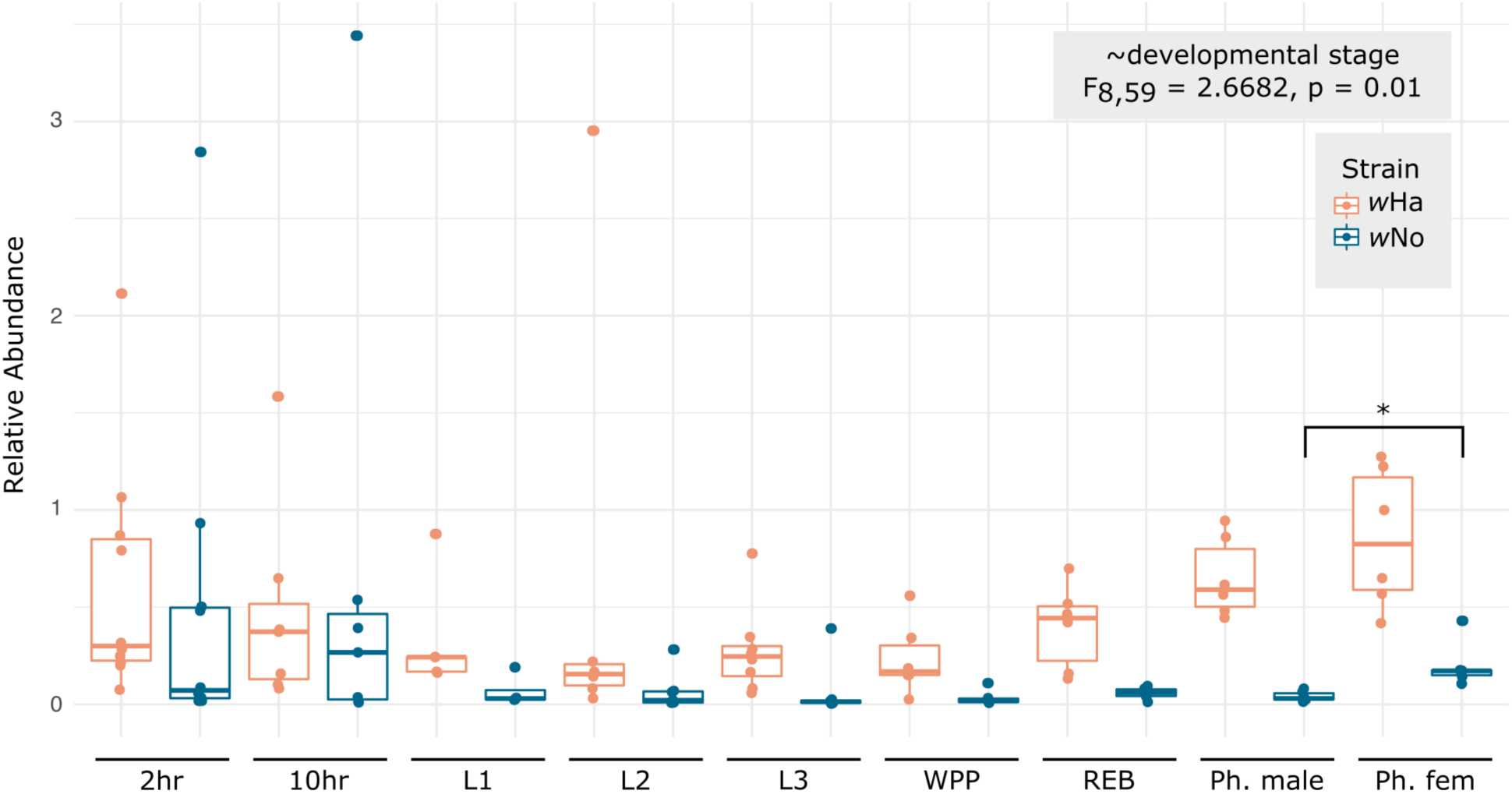
Coinfection is dynamic across development. Relative abundance of *w*Ha and *w*No across development. Developmental stages include, from left to right, 2-hour old embryos, 10-hour old embryos, 1st instar larvae (L1), 2nd instar larvae (L2), 3rd instar larvae (L3), white prepupae (WPP), red-eye bald pupal stage (REB), pharate (Ph.) males, and Ph. females.

### Transmission of the coinfection to embryos is strain-specific

The developmental series revealed that very young embryos had coinfections that were dissimilar to the infections in ovaries which raises questions about how the two *Wolbachia* are transmitted to the next generation (Figure 2B). However, the data presented in Figure 2B were generated from unmated females, so we sought to determine if the coinfection differed due to mating, which might explain why the embryos had differing ratios of the two *Wolbachia* strains. We found no significant difference in the coinfection between ovaries derived from three day-old mated and unmated females, and in both cases *w*No was significantly higher titer than *w*Ha (Figure 4A; ∼strain*mated status: F_1,12_ = 1.055, p = 0.3246; ∼mated status: F_1,12_ = 0.473, p = 0.5049; ∼strain: F_1,12_ = 22.891, p = 0.0005). We then used linear regression to assess the relationship between *w*Ha and *w*No in ovary and embryo samples with an eye towards the transmission dynamics. In both sample types there was a significant positive correlation between *w*Ha and *w*No, (ovaries: F_1,13_ = 45.13, p < 0.0001, r = 0.759; embryos: F_1,8_ = 133.9, p < 0.0001, r = 0.937). However, in ovaries *w*No was more than double the abundance of *w*Ha, whereas the two infections were closer to 1:1 in embryos (Figure 4B; ovaries: y=2.0281x+0.3804; embryos: y=1.3679x-0.4679). Therefore, transmission to embryos favors *w*Ha. This is also seen in the negative intercept along the y-axis (*w*No), indicating a higher likelihood that embryos might receive only *w*Ha, but not *w*No at especially low levels of overall transmission, even though ovaries contain double the titer of *w*No.

**Figure 4.**
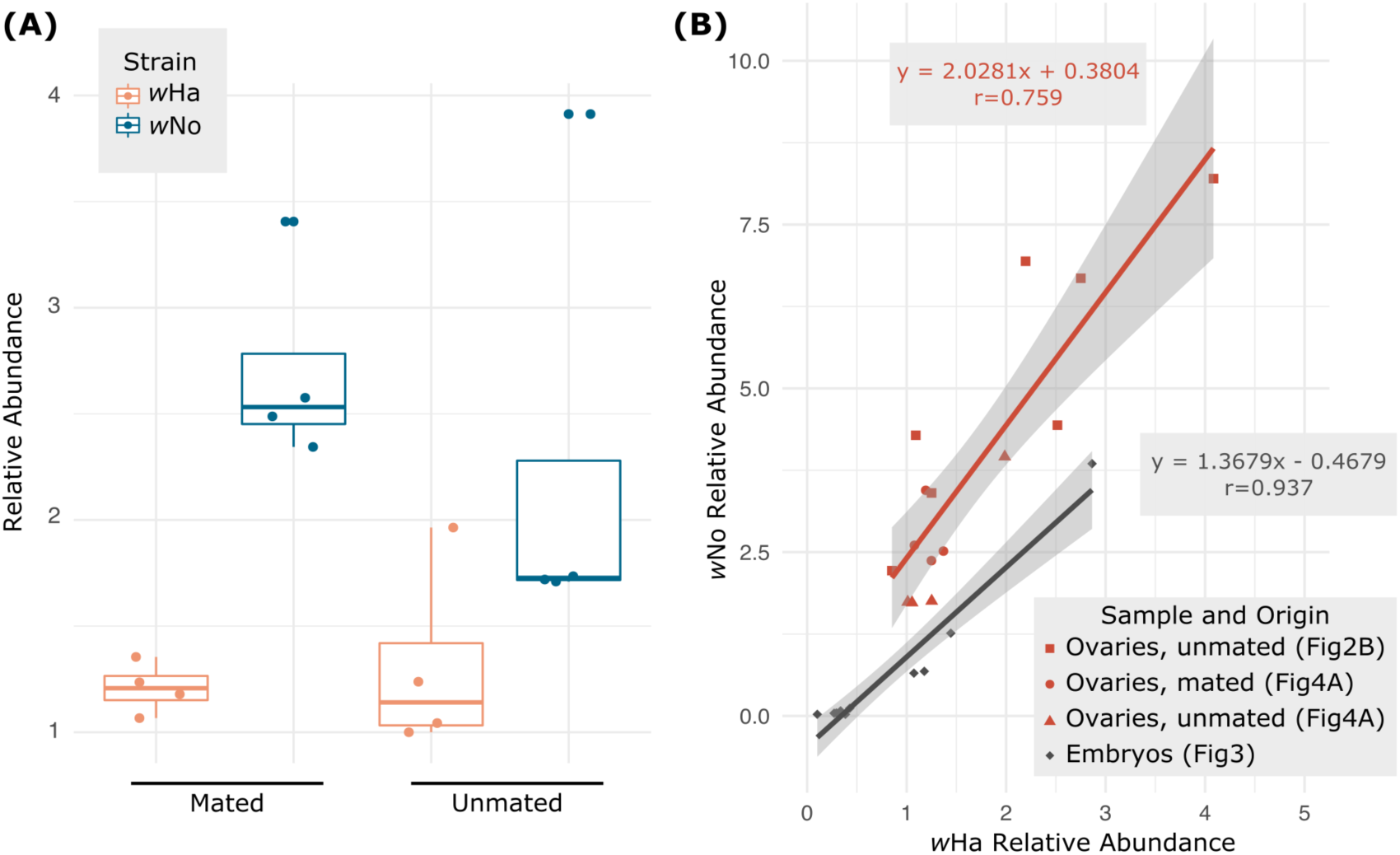
The ratio of *w*Ha and *w*No transmitted to embryos is not reflective of the coinfection in ovaries. **(A)** Titers of *w*Ha and *w*No do not significantly change upon mating. Newly eclosed females were collected and a subset were mated after 24-hours. Three days post eclosion, ovaries were dissected from the mated and unmated females. Only strain identity (*w*Ha versus *w*No) significantly affected titer (∼strain*mated status: F_1,12_ = 1.055, p = 0.3246; ∼mated status: F_1,12_ = 0.473, p = 0.5049; ∼strain: F_1,12_ = 22.891, p = 0.0005). **(B)** *w*Ha and *w*No titers are strongly correlated within ovaries, and within embryos. However, the ratios of *w*Ha:*w*No are significantly different between the two, indicated by the negative y-intercept (*w*No) for embryos as compared to ovaries.

### Heat stress facilitates destabilization of co-transmission

We hypothesized that we could perturb the transmission of the coinfection through a heat-mediated reduction in *Wolbachia* titers, which would facilitate a strong bottleneck and the opportunity to isolate individual *Wolbachia* strains. Indeed, subjecting flies to 30 °C for four days resulted in some F1 progeny (11.5%) that were lacking in one or both *Wolbachia* strains (Figure 5). This is in contrast to the offspring of flies reared at 25 °C, where the coinfection is stably transmitted: in our routine lab screens we have yet to find flies from this stock that do not carry both infections (n > 200).

**Figure 5.**
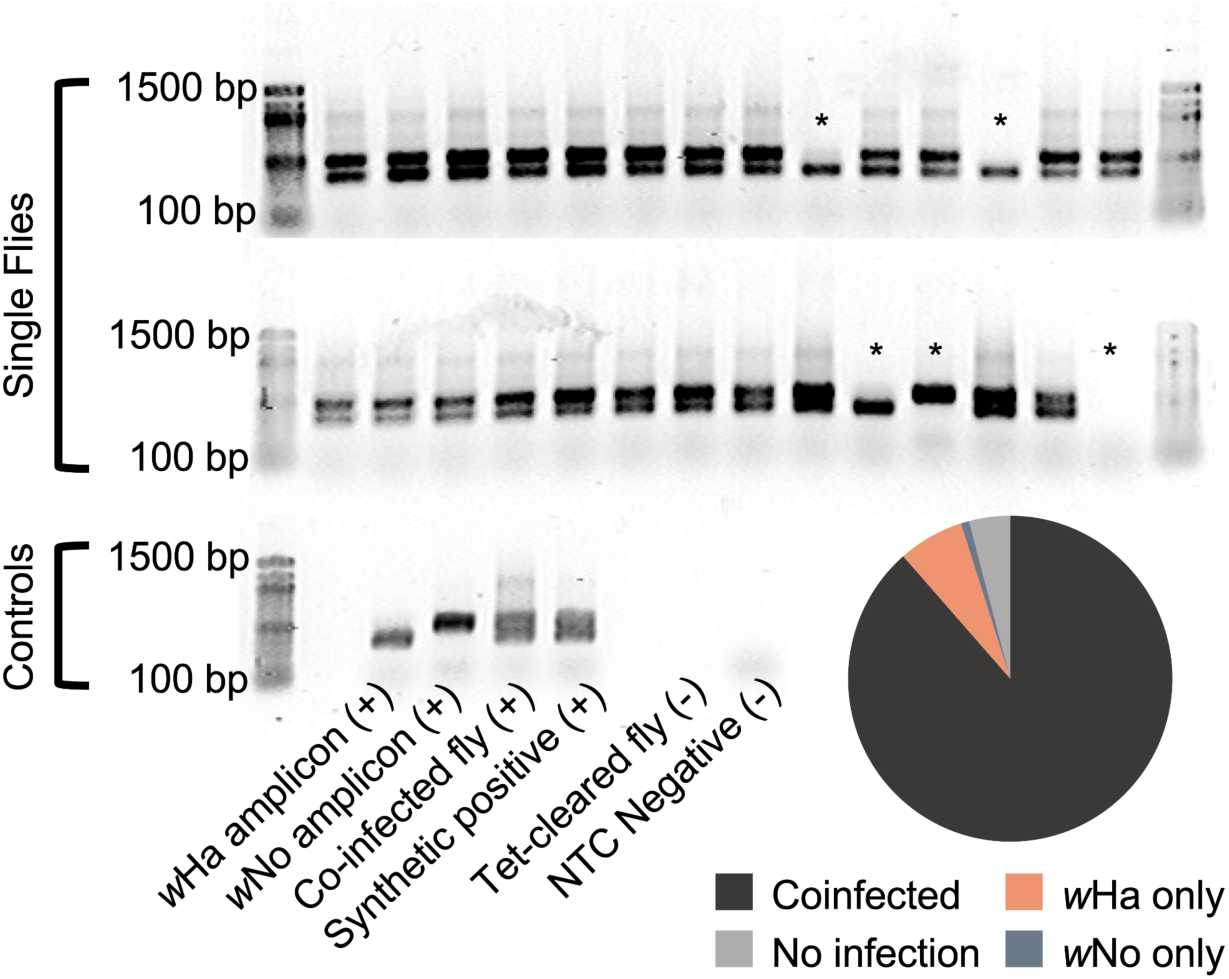
Heat stress destabilizes co-transmission of *w*Ha and *w*No. Gel electrophoresis of multiplex PCR assay indicating flies that have lost one or both *Wolbachia* infections (*). The “synthetic positive” control was generated by combining previously generated *w*Ha and *w*No amplicons in equimolar ratios. Negative controls include flies cleared of their *Wolbachia* infections, and no template controls (NTC). The pie chart summarizing the numbers of flies that lost *Wolbachia* infections (n = total 122 flies screened: *w*Ha only = 8, *w*No only = 1, uninfected = 5).

## DISCUSSION

We hypothesized that stability of multiple *Wolbachia* infections was made possible by some level of niche partitioning. That a coinfection is typically comprised of strains from different supergroups, with each supergroup having a unique set of clade-specific genes (49-51) supports this idea. In *w*Ha and *w*No we identified strain-specific proteins predicted to be involved in separate metabolic pathways, as well as proteins that may provide different mechanisms for host interaction and virulence. Indeed, *w*Ha and *w*No have different patterns of tissue tropism across males and females and show different transmission and growth dynamics across fly development.

While *w*Ha and *w*No titers differed significantly between the ovaries and early embryos, the mechanisms that resulted in differential transmission of *w*Ha and *w*No are still unclear. While *w*Ha and *w*No titers within the ovary are distinct from titers elsewhere in the body, there may be cell-type specificity within the ovary. Ovaries contain a variety of both somatic and germline cell-types, and there are documented examples of cell-type tropisms that also differ across *Wolbachia* strains (52, 53). Strain-specific imaging of whole ovarioles will allow us to determine how each *Wolbachia* strain is distributed within the ovary and in oocytes. The “assembly line” structure of *Drosophila* ovarioles offers a convenient way to capture changes in tissue specificity and titer that occur as eggs mature and may provide an explanation for the discrepancies in composition of the *Wolbachia* community that we see between whole ovaries and embryos.

After *w*Ha and *w*No are transmitted to the embryos, the coinfection seems to converge on a pattern consisting of a relatively low and stable population of *w*No and a comparatively high level of *w*Ha that persists throughout development. When the adults emerge, we see the first evidence of increasing *w*No titers in females. Our data suggest that the switch from the high *w*Ha:*w*No ratio seen in juveniles to the relatively equal *w*Ha:*w*No titers of three day old females occurs during adulthood, not metamorphosis. This process may be linked to ovary maturation as an adult rather than imaginal disc differentiation during the pupal period, but more in-depth analyses of the imaginal discs and the adult female maturation period are needed to tease this apart.

The differences in infection between ovaries and testes raise several questions about the reproductive manipulation induced by these strains: Cytoplasmic Incompatibility (CI). In the testes, CI results in altered sperm that cause embryonic lethality, unless “rescued” by a complementary infection in the oocyte (54). In the case of coinfections, typically each strain-specific alteration of the sperm requires a matching rescue or antidote in the embryo (10, 55), and previous studies indicate that *w*Ha and *w*No are not fully capable of rescuing the other strain’s CI induction (46). These CI induction and rescue processes are mediated by *Wolbachia* “Cif” proteins, and there is strong evidence that the level of Cif expression, and the availability of strain-specific cognate partners is critical for proper induction and rescue (54, 56-59). Given this, it was interesting to find that the ratio of *w*Ha to *w*No within the testes was more variable between individuals than it was across ovaries (in which *w*Ha and *w*No titers were strongly correlated). Additionally, *w*Ha was the dominant strain in testes, as compared to *w*No being dominant in the ovaries. It is not clear if the ratios of *w*Ha and *w*No infections in the gonad tissues are reflective of the level of Cif proteins in gametes, and ultimately the level of induction and rescue caused by each strain. Perhaps expression and deposition of Cif proteins is regulated in a cell-type-specific or co-infection sensitive manner. Finally, we do not know if CI rescue is oocyte-autonomous, or if Cif proteins are transported between cell types (e.g., from somatic follicle cells to the oocyte). Which cell types do *Wolbachia* need to be in, and at what time points in gametogenesis in order to cause or rescue CI? Perhaps the quantity of Cif proteins from each strain that are deposited in spermatozoa and oocytes are tightly regulated such that they more closely mirror each other. A combination of molecular approaches to assess Cif protein abundance in gametes, combined with genetic tools to test for cell autonomy will be useful for understanding these processes, and ultimately how CI is regulated.

Finally, we demonstrated that heat stress disrupts vertical transmission of *w*Ha and *w*No through an unknown mechanism. We hypothesize that heat stress negatively impacts *Wolbachia* titers (60), causing the bacteria to be “diluted” as cells in the ovary chain divide. In rare instances, a developing oocyte will receive *Wolbachia* of only one strain or no *Wolbachia* at all. Using a heat treatment, we recovered more flies that only had the *w*Ha strain (and had lost *w*No), and only one example of a fly that only had *w*No (n = 1). This may be due to the preferential transmission of *w*Ha that we saw when comparing ovary and embryo coinfections, or potentially strain-specific differences in heat-sensitivity. Indeed, a recent study showed that temperature is a strong driver of *Wolbachia* transmission and spread at large scales (61), and there are many other examples of high temperatures that result in full or partial cures of *Wolbachia* (60). Our ability to segregate the strains into mono-infections in the same genomic background will be a useful tool for exploring the strain-specific contributions to host physiology, and for understanding the interactions between coinfecting *Wolbachia*. Indeed, a combination of factors likely governs *Wolbachia* community dynamics, and it is unclear if *w*Ha and *w*No interactions with each other are competitive, synergistic, or perhaps parasitic. Disentangling the relative contributions of each strain to the stability of the coinfection will inform efforts to establish multiple infections of selected symbionts and contribute to understanding the dynamics of the intracellular community more broadly.

## DECLARATIONS

## Acknowledgements

This work was supported by UMN AGREETT startup funds to ARIL. Many thanks to Brandon Cooper for gifting us the *Drosophila simulans* stock. LCF was supported by UMN DOVE and UMN CFANS Match fellowships. MWJ was supported by an Excellence in Entomology Fellowship from UMN.

## Data Availability

**Supplemental File S1**. Contains supplemental figures.

**Figure S1**. Within-fly gonad and carcass infection dynamics.

**Figure S2**. *w*Ha and *w*No correlation across development.

**Supplemental File S2:** Contains supplemental tables. Metadata are in the first tab of the file.

**Table S1**. FtsZ protein accession numbers used for phylogenetic reconstruction.

**Table S2**. KEGG annotations for *w*Ha and *w*No specific proteins.

**Tables S3-S7**. qPCR data by figure.

## REFERENCES

1. Duron O, Bouchon D, Boutin S, Bellamy L, Zhou L, Engelstädter J, Hurst GD. 2008. The diversity of reproductive parasites among arthropods: Wolbachia do not walk alone. BMC Biol 6:1–12.

2. Duron O, Hurst GD. 2013. Arthropods and inherited bacteria: from counting the symbionts to understanding how symbionts count. BMC Biol 11:1–4.

3. Dumler JS, Barbet AF, Bekker C, Dasch GA, Palmer GH, Ray SC, Rikihisa Y, Rurangirwa FR. 2001. Reorganization of genera in the families Rickettsiaceae and Anaplasmataceae in the order Rickettsiales: unification of some species of Ehrlichia with Anaplasma, Cowdria with Ehrlichia and Ehrlichia with Neorickettsia, descriptions of six new species combinations and designation of Ehrlichia equi and’HGE agent’as subjective synonyms of Ehrlichia phagocytophila. Int J Syst Evol Microbiol 51:2145–2165.

4. Werren JH, Baldo L, Clark ME. 2008. Wolbachia: master manipulators of invertebrate biology. Nat Rev Micro 6:741–751.

5. Kaur R, Shropshire JD, Cross KL, Leigh B, Mansueto AJ, Stewart V, Bordenstein SR, Bordenstein SR. 2021. Living in the endosymbiotic world of Wolbachia: A centennial review. Cell Host & Microbe.

6. Werren JH, Windsor D, Guo LR. 1995. Distribution of Wolbachia among neotropical arthropods. Proc R Soc Lond B 262:197–204.

7. Bordenstein SR, Werren JH. 1998. Effects of A and B Wolbachia and host genotype on interspecies cytoplasmic incompatibility in Nasonia. Genetics 148:1833–1844.

8. Ellegaard KM, Klasson L, Naslund K, Bourtzis K, Andersson SGE. 2013. Comparative genomics of Wolbachia and the bacterial species concept. PLoS Genet 9:e1003381.

9. Dutton TJ, Sinkins SP. 2004. Strain-specific quantification of Wolbachia density in Aedes albopictus and effects of larval rearing conditions. Insect Mol Biol 13:317–322.

10. Dobson S, Rattanadechakul W, Marsland E. 2004. Fitness advantage and cytoplasmic incompatibility in Wolbachia single-and superinfected Aedes albopictus. Heredity 93:135–142.

11. Bordenstein SR, O’hara FP, Werren JH. 2001. Wolbachia-induced incompatibility precedes other hybrid incompatibilities in Nasonia. Nature 409:707–710.

12. Merçot H, Poinsot D. 1998. Wolbachia transmission in a naturally bi-infected Drosophila simulans strain from New-Caledonia. Entomol Exp Appl 86:97–103.

13. Kent BN, Salichos L, Gibbons JG, Rokas A, Newton ILG, Clark ME, Bordenstein SR. 2011. Complete bacteriophage transfer in a bacterial endosymbiont (Wolbachia) determined by targeted genome capture. Genome Biol Evol 3:209–218.

14. Chafee ME, Funk DJ, Harrison RG, Bordenstein SR. 2010. Lateral phage transfer in obligate intracellular bacteria (Wolbachia): verification from natural populations. Mol Biol Evol 27:501–5.

15. Baldo L, Bordenstein S, Wernegreen JJ, Werren JH. 2006. Widespread recombination throughout Wolbachia genomes. Mol Biol Evol 23:437–449.

16. Lindsey ARI, Bhattacharya T, Newton ILG, Hardy RW. 2018. Conflict in the intracellular lives of endosymbionts and viruses: A mechanistic look at Wolbachia-mediated pathogen-blocking. Viruses 10:141.

17. Joubert DA, Walker T, Carrington LB, De Bruyne JT, Kien DHT, Hoang NLT, Chau NVV, Iturbe-Ormaetxe I, Simmons CP, O’Neill SL. 2016. Establishment of a Wolbachia superinfection in Aedes aegypti mosquitoes as a potential approach for future resistance management. PLoS Path 12:e1005434.

18. Ross PA, Turelli M, Hoffmann AA. 2019. Evolutionary ecology of Wolbachia releases for disease control. Annu Rev Genet 53:93–116.

19. Walker T, Johnson PH, Moreira LA, Iturbe-Ormaetxe I, Frentiu FD, McMeniman CJ, Leong YS, Dong Y, Axford J, Kriesner P, Lloyd AL, Ritchie SA, O’Neill SL, Hoffmann AA. 2011. The wMel Wolbachia strain blocks dengue and invades caged Aedes aegypti populations. Nature 476:450–U101.

20. Hoffmann AA, Montgomery BL, Popovici J, Iturbe-Ormaetxe I, Johnson PH, Muzzi F, Greenfield M, Durkan M, Leong YS, Dong Y, Cook H, Axford J, Callahan AG, Kenny N, Omodei C, McGraw EA, Ryan PA, Ritchie SA, Turelli M, O’Neill SL. 2011. Successful establishment of Wolbachia in Aedes populations to suppress dengue transmission. Nature 476:454–U107.

21. Hoffmann AA, Ross PA, Rasic G. 2015. Wolbachia strains for disease control: ecological and evolutionary considerations. Ecol Evol 8:751–68.

22. Black C, Bermudez L, Young L, Remington J. 1990. Co-infection of macrophages modulates interferon gamma and tumor necrosis factor-induced activation against intracellular pathogens. The Journal of Experimental Medicine 172:977–980.

23. Erickson AK, Jesudhasan PR, Mayer MJ, Narbad A, Winter SE, Pfeiffer JK. 2018. Bacteria facilitate enteric virus co-infection of mammalian cells and promote genetic recombination. Cell Host & Microbe 23:77-88. e5.

24. Nakamura S, Davis KM, Weiser JN. 2011. Synergistic stimulation of type I interferons during influenza virus coinfection promotes Streptococcus pneumoniae colonization in mice. J Clin Invest 121.

25. Spier A, Stavru F, Cossart P. 2019. Interaction between intracellular bacterial pathogens and host cell mitochondria. Bacteria and Intracellularity:1–13.

26. Heddi A, Grenier A-M, Khatchadourian C, Charles H, Nardon P. 1999. Four intracellular genomes direct weevil biology: nuclear, mitochondrial, principal endosymbiont, and Wolbachia. Proc Natl Acad Sci 96:6814–6819.

27. Sassera D, Beninati T, Bandi C, Bouman EA, Sacchi L, Fabbi M, Lo N. 2006. ‘Candidatus Midichloria mitochondrii’, an endosymbiont of the tick Ixodes ricinus with a unique intramitochondrial lifestyle. Int J Syst Evol Microbiol 56:2535–2540.

28. Ramírez-Puebla ST, Ormeño-Orrillo E, Vera-Ponce de León A, Lozano L, Sanchez-Flores A, Rosenblueth M, Martínez-Romero E. 2016. Genomes of Candidatus Wolbachia bourtzisii w DacA and Candidatus Wolbachia pipientis w DacB from the Cochineal Insect Dactylopius coccus (Hemiptera: Dactylopiidae). G3: Genes, Genomes, Genetics 6:3343–3349.

29. Liang X, Liu J, Bian G, Xi Z. 2020. Wolbachia inter-strain competition and inhibition of expression of cytoplasmic incompatibility in mosquito. Frontiers in Microbiology 11:1638.

30. Kang L, Ma X, Cai L, Liao S, Sun L, Zhu H, Chen X, Shen D, Zhao S, Li C. 2003. Superinfection of Laodelphax striatellus with Wolbachia from Drosophila simulans. Heredity 90:71–76.

31. Rousset F, Braig HR, O’Neill SL. 1999. A stable triple Wolbachia infection in Drosophila with nearly additive incompatibility effects. Heredity 82:620–627.

32. Walker T, Song S, Sinkins SP. 2009. Wolbachia in the Culex pipiens group mosquitoes: introgression and superinfection. J Hered 100:192–196.

33. Ant TH, Sinkins SP. 2018. A Wolbachia triple-strain infection generates self-incompatibility in Aedes albopictus and transmission instability in Aedes aegypti. Parasites & Vectors 11:1–7.

34. Schneider DI, Riegler M, Arthofer W, Merçot H, Stauffer C, Miller WJ. 2013. Uncovering Wolbachia diversity upon artificial host transfer. PLoS One 8:e82402.

35. Fu Y, Gavotte L, Mercer DR, Dobson SL. 2010. Artificial triple Wolbachia infection in Aedes albopictus yields a new pattern of unidirectional cytoplasmic incompatibility. Appl Environ Microbiol 76:5887–5891.

36. Osborne SE, Leong YS, O’Neill SL, Johnson KN. 2009. Variation in antiviral protection mediated by different Wolbachia strains in Drosophila simulans. PLoS Path 5:e1000656.

37. Osborne SE, Iturbe-Ormaetxe I, Brownlie JC, O’Neill SL, Johnson KN. 2012. Antiviral protection and the importance of Wolbachia density and tissue tropism in Drosophila simulans. Appl Environ Microbiol 78:6922–6929.

38. Poinsot D, Montchamp-Moreau C, Merçot H. 2000. Wolbachia segregation rate in Drosophila simulans naturally bi-infected cytoplasmic lineages. Heredity 85:191–198.

39. Mouton L, Dedeine F, Henri H, Boulétreau M, Profizi N, Vavre F. 2004. Virulence, multiple infections and regulation of symbiotic population in the Wolbachia-Asobara tabida symbiosis. Genetics 168:181–189.

40. Mouton L, Henri H, Bouletreau M, Vavre F. 2003. Strain-specific regulation of intracellular Wolbachia density in multiply infected insects. Mol Ecol 12:3459–3465.

41. Tatusova T, DiCuccio M, Badretdin A, Chetvernin V, Nawrocki EP, Zaslavsky L, Lomsadze A, Pruitt KD, Borodovsky M, Ostell J. 2016. NCBI prokaryotic genome annotation pipeline. Nucleic Acids Res 44:6614–6624.

42. Lechner M, Findeiß S, Steiner L, Marz M, Stadler PF, Prohaska SJ. 2011. Proteinortho: detection of (co-) orthologs in large-scale analysis. BMC Bioinformatics 12:1–9.

43. Kanehisa M, Sato Y, Morishima K. 2016. BlastKOALA and GhostKOALA: KEGG tools for functional characterization of genome and metagenome sequences. J Mol Biol 428:726–731.

44. Katoh K, Rozewicki J, Yamada KD. 2019. MAFFT online service: multiple sequence alignment, interactive sequence choice and visualization. Briefings in Bioinformatics 20:1160–1166.

45. Bhattacharya T, Newton ILG, Hardy RW. 2017. Wolbachia elevates host methyltransferase expression to block an RNA virus early during infection. PLoS Path 13:e1006427.

46. James A, Dean M, McMahon M, Ballard J. 2002. Dynamics of double and single Wolbachia infections in Drosophila simulans from New Caledonia. Heredity 88:182–189.

47. R Core Team. 2014. R: A language and environment for statistical computing, URL http://www.R-project.org/, R Foundation for Statistical Computing, Vienna, Austria.

48. Oksanen J, Blanchet FG, Kindt R, Legendre P, Minchin PR, O’Hara R, Simpson GL, Solymos P, Stevens M, Wagner H. 2015. vegan: Community Ecology Package. R package version 2.0-1. http://CRANR-projectorg/package=vegan.

49. Lindsey ARI, Werren JH, Richards S, Stouthamer R. 2016. Comparative genomics of a parthenogenesis-inducing Wolbachia symbiont. G3: Genes|Genomes|Genetics 6:2113–2123.

50. Lindsey ARI. 2020. Sensing, Signaling, and Secretion: A review and analysis of systems for regulating host interaction in Wolbachia. Genes 11:813.

51. Rice DW, Sheehan KB, Newton IL. 2017. Large-scale identification of Wolbachia pipientis effectors. Genome Biol Evol 9:1925–1937.

52. Fast EM, Toomey ME, Panaram K, Desjardins D, Kolaczyk ED, Frydman HM. 2011. Wolbachia enhance Drosophila stem cell proliferation and target the germline stem cell niche. Science 334:990–992.

53. Frydman HM, Li JM, Robson DN, Wieschaus E. 2006. Somatic stem cell niche tropism in Wolbachia. Nature 441:509–12.

54. Beckmann JF, Bonneau M, Chen H, Hochstrasser M, Poinsot D, Merçot H, Weill M, Sicard M, Charlat S. 2019. The toxin–antidote model of cytoplasmic incompatibility: genetics and evolutionary implications. Trends Genet.

55. Sinkins S, Braig H, O’Neill SL. 1995. Wolbachia superinfections and the expression of cytoplasmic incompatibility. Proceedings of the Royal Society of London Series B: Biological Sciences 261:325–330.

56. Shropshire JD, Bordenstein SR. 2019. Two-By-One model of cytoplasmic incompatibility: Synthetic recapitulation by transgenic expression of cifA and cifB in Drosophila. PLoS Genet 15.

57. Chen H, Ronau JA, Beckmann JF, Hochstrasser M. 2019. A Wolbachia nuclease and its binding partner provide a distinct mechanism for cytoplasmic incompatibility. Proc Natl Acad Sci 116:22314–22321.

58. Lindsey ARI, Rice DW, Bordenstein SR, Brooks AW, Bordenstein SR, Newton ILG. 2018. Evolutionary genetics of cytoplasmic incompatibility genes cifA and cifB in prophage WO of Wolbachia. Genome Biol Evol 10:434–451.

59. Beckmann JF, Ronau JA, Hochstrasser M. 2017. A Wolbachia deubiquitylating enzyme induces cytoplasmic incompatibility. Nat Micro 2:17007.

60. López-Madrigal S, Duarte EH. 2019. Titer regulation in arthropod-Wolbachia symbioses. FEMS Microbiol Lett 366:fnz232.

61. Hague MT, Shropshire JD, Caldwell CN, Statz JP, Stanek KA, Conner WR, Cooper BS. 2022. Temperature effects on cellular host-microbe interactions explain continent-wide endosymbiont prevalence. Curr Biol 32:878-888. e8.

